# Fluorescent nanodiamond phantoms for spatial resolution benchmarking in two- and three-photon microscopy

**DOI:** 10.64898/2026.01.12.699010

**Authors:** Filip Janiak, Michael Forsthofer, Jakub Czubek, Sinzi Pop, Yangting Fu, Lucia Prieto-Godino, Hans-Ulrich Fried, Małgorzata Szczerska, Tom Baden

## Abstract

Reliable spatial-resolution benchmarking is essential for multiphoton microscopy, yet conventional fluorescent-bead samples are imperfect references for biological imaging. Their fluorescence and sample properties can vary with preparation, storage and excitation conditions, while the surrounding agarose does not reproduce the depth-dependent optical degradation encountered in tissue. Here, we evaluate fluorescent nanodiamond phantoms as a more robust alternative for two- and three-photon microscopy. Nanodiamonds provide well-defined point-spread-function measurements across different microscope configurations, can be repeatedly imaged over long periods using the same physical sample, and remain usable across a broad range of excitation powers. Crucially, their optically heterogeneous matrix produces progressive broadening of the measured PSF with imaging depth, capturing a defining feature of real biological imaging that conventional bead phantoms largely miss. Nanodiamond phantoms therefore provide a durable and experimentally forgiving reference for benchmarking not only nominal spatial resolution, but how that resolution is maintained as imaging conditions become more challenging.

## Introduction

Two- and three-photon microscopy (2PM and 3PM) are nonlinear imaging techniques that excite fluorophores through the near-simultaneous absorption of two or three infrared photons, respectively. By confining excitation to a small focal volume, these techniques enable optical sectioning with reduced out-of-focus excitation^1,2^. Two-photon microscopy is widely used for *in vivo* imaging of neuronal activity and fine neuronal structures across brain regions and species^3–8^.

Accurate spatial resolution measurements ensure that fine structures are correctly and reproducibly visualized. Accordingly, robust methods for spatial-resolution assessment are essential for meaningful within- and cross-platform comparisons^9^.

The point spread function (PSF) is a fundamental descriptor of a microscope’s spatial resolution^9,10^. It represents the three-dimensional response of the imaging system to a point source, effectively characterizing how light from a single point object spreads in the lateral (x, y) and axial (z) dimensions. This makes the PSF a central metric for validating system performance, optimizing imaging parameters, ensuring reproducibility across experiments, and comparing different multiphoton systems.

To accurately measure the PSFs of a multiphoton microscope, researchers typically image sub-resolution fluorescent beads (ideally 100–200 nm in diameter) embedded in low-melting-point agarose gel^9^. A three-dimensional image stack of an isolated bead is acquired with adequate voxel sampling to resolve the PSF. Full width at half maximum (FWHM) measurements are taken along the lateral and axial dimensions to estimate spatial resolution^10,11^. To identify field-dependent variation or optical distortion, measurements are repeated across multiple regions of the sample. This approach provides a practical baseline for assessing system performance^12^. However, sample preparation and local aberrations can introduce variability in the measured PSF^13,14^. Further variability can stem from bead immobilization and storage conditions^9^. Variations in bead distribution and photobleaching can also affect repeated measurements^9^. Depending on the fluorophore and storage conditions, long-term storage may also affect probe performance^15^. In addition, refractive-index mismatch introduced by the sample mounting geometry and coverslip can generate spherical aberrations^14^.

Two-photon microscopy is now widely used, while three-photon microscopy extends nonlinear imaging to greater depths. Its different excitation regime and the effects of scattering and aberrations in deep tissue place additional demands on calibration and performance assessment^16–18^. Fluorescent beads remain widely used for three-photon PSF characterisation, including at substantial imaging depths^19^. A practical reference material should therefore remain detectable and support lateral and axial PSF measurements across imaging depths, excitation powers and microscope platforms. Adaptive-optics three-photon microscopy further demonstrates the importance of aberration correction for high-resolution imaging deep in the brain^20^. Fluorescent nanodiamonds have also been investigated as optical probes in cells, including the detection of individual internalised nanoparticles^21^.

Fluorescent nanodiamonds have previously been proposed as photostable calibration targets for nonlinear microscopy and longitudinal instrument quality control^22^. Here, we extend this concept to three-dimensional, optically heterogeneous phantoms for PSF measurements under both two- and three-photon excitation. Beyond their bright and stable fluorescence under nonlinear excitation^22,23^, nanodiamonds provide key practical advantages for PSF characterization. Their intrinsic fluorescence is highly resistant to photobleaching and their exceptional chemical stability enables long-term preservation^24,25^. The samples used in this study were prepared in 2018 and remained suitable for PSF measurements in 2025 after long-term storage at room temperature; their optical and mechanical properties were not quantified at both time points. Reusing the same phantom across imaging sessions avoids the need to prepare a new sample for each measurement. Moreover, the tested formulations show refractive indices within the range reported for some biological tissues, providing an optically heterogeneous environment for benchmarking imaging performance. Together with their reported fluorescence stability^24,26^, these features motivate the evaluation of nanodiamond-based phantoms as reusable reference materials for PSF measurements in two- and three-photon microscopy.

## Results

Nanodiamonds are crystalline carbon nanoparticles produced by high-pressure or detonation synthesis^26,27^. Their reported fluorescence and resistance to photobleaching have motivated their use as optical reference probes^23,25^. In this study, we assessed stored nanodiamond-containing phantoms for PSF measurement; photobleaching and biocompatibility were not measured directly.

### Nanodiamond phantoms as calibration materials

We used nanodiamond phantoms originally developed to approximate selected physical properties of human tissue in the context of medical imaging (calibrating MRI scans)^28^. The samples were prepared using an agar–carrageenan gel matrix in distilled water. Two nanodiamond series were prepared: the nano series containing 8–12% (v/v) of an ND-standard suspension in DMSO (Adámas Nanotechnologies, 5 nm particles), and the dnp series containing 8–12% (v/v) of a hydrophilic RT-DND-L suspension (Ray Techniques Ltd., 4–5 nm particles). The heated gel base was combined with the prepared nanodiamond suspensions; after mixing, the mixtures underwent a further autoclaving step before being poured into moulds and allowed to solidify (see Methods for details). DMSO has previously been investigated as a dispersion medium for detonation nanodiamonds^29^.

Figure 1a presents scanning electron micrographs (SEM) of an 8% dnp phantom. The images reveal a heterogeneous distribution of nanodiamond structures, including aggregates below 200 nm, comparable in size to standard fluorescent beads used for PSF calibration. Individual nanodiamond particles are nominally 4–5 nm in diameter. Previous dynamic light scattering (DLS) measurements of the RT-DND suspension used in the phantom formulation reported an average aggregate size of approximately 70–80 nm^28^. This value refers to aggregates in the precursor suspension, not to a measured size distribution of fluorescent emitters selected within the finished gel.

**Figure 1.**
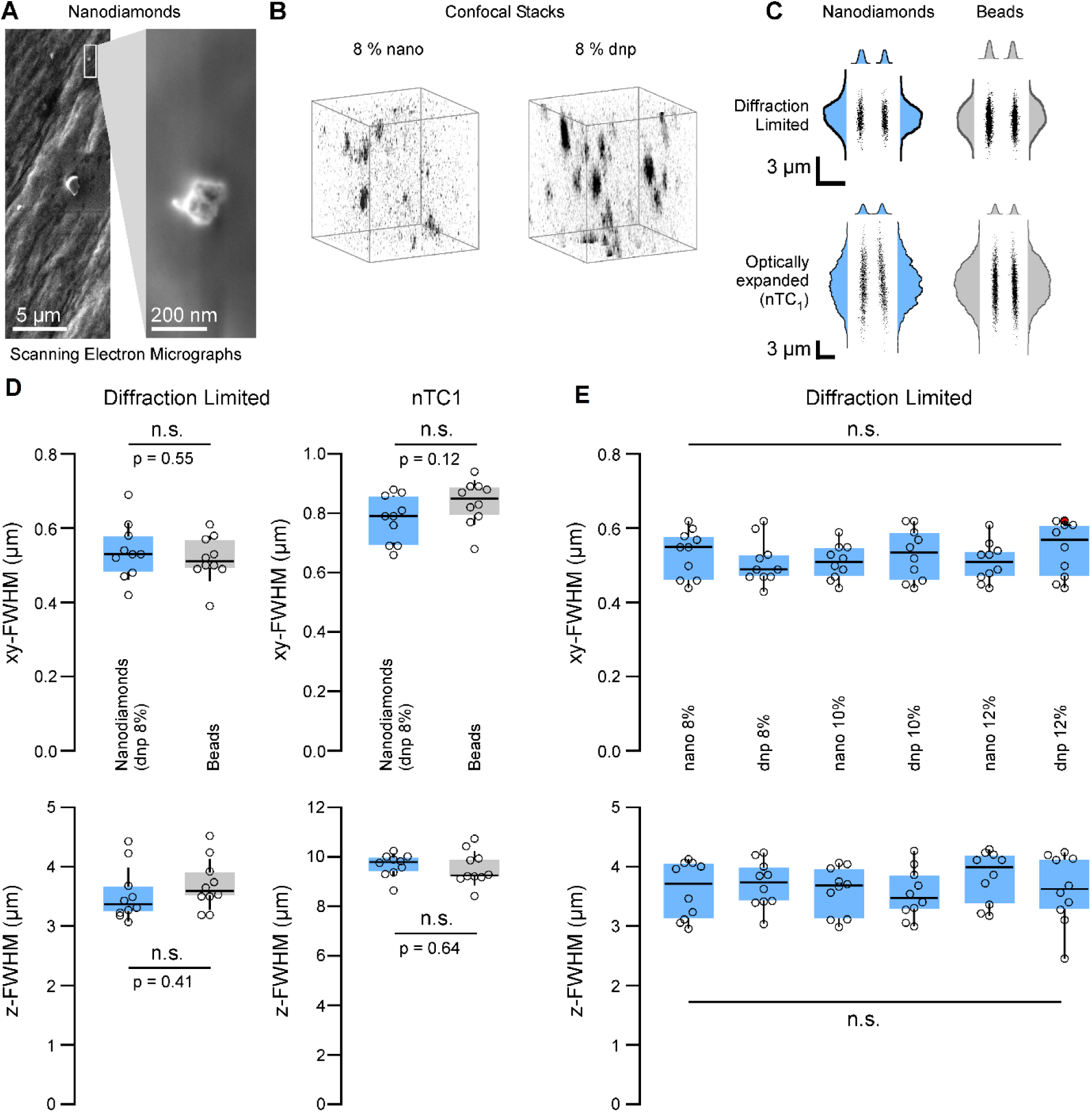
Structural characterisation and two-photon PSF measurements of nanodiamond-based phantoms. (a) Scanning electron microscopy (SEM) images of an 8% dnp nanodiamond phantom reveal heterogeneous nanodiamond aggregates with a substantial fraction below 200 nm, comparable in size to standard fluorescent beads used for PSF calibration. (b) Volumetric confocal microscopy of nano 8% and dnp 8% phantoms under 561-nm excitation shows bright punctate fluorescent structures distributed throughout the gel, together with diffuse background luminescence. The puncta are consistent with fluorescent nanodiamond aggregates, but no matrix-only or emission-spectral control was performed to determine the origin of all detected signal. (c) Representative xz and yz cross-sections of two-photon PSFs recorded from nanodiamonds (dnp 8%) and commercial fluorescent beads (Invitrogen 505/515) under diffraction-limited (DL) and non-telecentric (nTC) imaging configurations. All PSFs were acquired using 920 nm excitation at 3 mW on the same two-photon system. (d) No statistically significant nanodiamond–bead differences in lateral or axial FWHM were detected in either DL or nTC mode under the tested conditions; these comparisons do not constitute a formal test of equivalence. (e) PSF measurements obtained from all six nanodiamond phantom formulations (DL configuration) show no statistically significant FWHM differences among the tested formulations under the reported conditions; batch-to-batch reproducibility was not assessed. For quantitative panels (d,e), open circles indicate individual emitter measurements; the boxes extend from the first to the third quartile (Tukey method), centre lines indicate the median, and whiskers extend from the mean − 1 SD to the mean + 1 SD.

Confocal z-stacks acquired under 561-nm excitation revealed bright, punctate fluorescent emitters distributed throughout both nanodiamond phantom formulations, together with diffuse background luminescence within the gel (Figure 1b). The punctate fluorescence is consistent with emission from nanodiamond aggregates embedded in the matrix. No fluorescent dye was added to the formulation, but neither a matrix-only control nor emission-spectral measurements were performed; thus, these images do not independently establish the origin of each punctum or of the diffuse background. The isolated bright structures nevertheless provided suitable targets for PSF measurements.

### Nanodiamond phantoms enable robust and reproducible two-photon PSF measurements

We demonstrate that nanodiamonds give usable PSFs under 2-photon excitation. We compared two-photon PSF measurements (920 nm excitation, 3 mW at the sample, Coherent Vision S femtosecond laser) from one of the nanodiamond samples (8% dnp) with those from a standard fluorescent bead sample (Invitrogen 505/515, P7220) using two-photon microscope Setup A (University of Sussex). Representative *xz* and *yz* cross-sections were extracted from *three-dimensional* image stacks acquired sequentially on the same imaging system configured in two modes: diffraction-limited (DL) and an experimentally expanded PSF generated with non-telecentric (nTC) optics^30^ (Figure 1c). In both configurations, no statistically significant nanodiamond–bead difference in PSF width was detected (Figure 1d), supporting the use of nanodiamonds for PSF measurements under the tested conditions.

Finally, Figure 1e compares PSF measurements obtained from all six phantom types under identical two-photon microscopy conditions (DL configuration). No statistically significant difference in the measured FWHM was detected among the six tested formulations in this experiment; this does not establish formal equivalence or batch-to-batch reproducibility.

### Optical properties and depth-dependent PSF measurements in nanodiamond phantoms

To assess PSF measurements in an optically heterogeneous environment, we compared phantom formulations containing different nanodiamond suspensions and concentrations. We first compared the appearance of the specimens with the conventional agarose-bead reference and then measured their refractive indices. These observations identify differences among the tested formulations, although controlled, independently reproducible tuning of refractive index or scattering was not established.

Figure 2a shows representative photographs of the various samples used in this study: an agarose-bead phantom, the nanodiamond-based phantoms (nano 8% and dnp 8%) and the exposed surface of an adult mouse brain viewed through a cranial window. The nanodiamond-based phantoms appear more optically heterogeneous than the uniform agarose-bead sample. This qualitative photographic comparison does not quantify their similarity to brain-tissue scattering.

**Figure 2.**
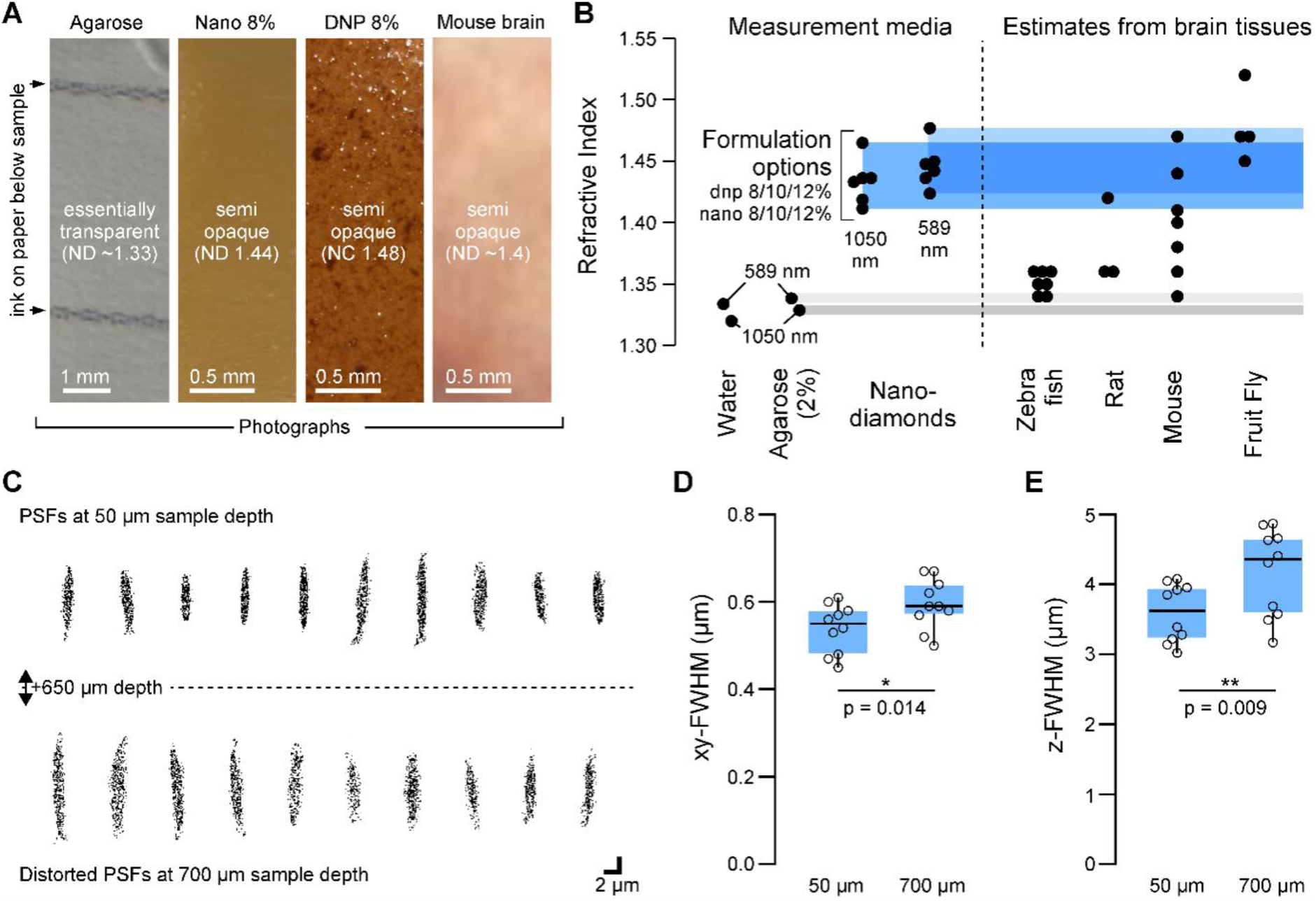
Optical properties, tissue relevance, and depth-dependent PSF performance of nanodiamond phantoms. (a) Representative photographs of the different sample types: an agarose-bead phantom, nanodiamond phantoms (nano 8% and dnp 8%), and the exposed surface of the adult mouse brain viewed through a cranial window. The nanodiamond phantoms show greater visible opacity and structural heterogeneity than the homogeneous agarose-bead sample, qualitatively resembling the appearance of brain tissue. (b) Refractive indices (RI) of distilled water, agarose gel, and six nanodiamond-based formulations measured at 589 nm and 1050 nm, compared with RI estimates from commonly used neuroscience model organisms (mouse, rat, zebrafish, drosophila). Several phantom RI values fall within reported brain-tissue ranges; literature estimates may have been acquired under different conditions, and phantom RI was not measured at 1300 nm. (c) Two-photon PSFs recorded in a nano 10% phantom at shallow (0–50 µm) and deep (∼700 µm) imaging depths under identical optical conditions. Representative xz/yz sections illustrate measured broadening with depth, consistent with increased optical perturbations within the phantom. (d) Lateral xy-FWHM values increase significantly between approximately 50 µm and 700 µm imaging depth (p = 0.014). (e) Axial z-FWHM values show a corresponding increase with depth (p = 0.009). For quantitative panels (d,e), open circles indicate individual measurements; boxes show Q1–Q3 (Tukey quartiles), centre lines show the median, and whiskers span the mean ± 1 SD.

Figure 2b summarises the refractive indices (RI) measured directly for distilled water, 2% agarose gel prepared with water, and the six nanodiamond phantoms used in this study (nano and dnp series; 8%, 10% and 12% v/v). Both nanodiamond series were measured at 589 and 1050 nm. For comparison, these data are plotted alongside published RI estimates for brain tissues of animal models commonly used in neuroscience: mouse^31–34^, rat^33,35^, zebrafish^36–40^ and fruit fly^41–43^. Several phantom values fall within the range reported for some animal brains, whereas the RIs of water and agarose are generally lower. The literature values may have been acquired under different conditions; the RI comparison does not establish matched tissue scattering.

In addition to the variation in RI across different tissue types, it is important to note that the refractive index measurements were collected at two wavelengths, 589 nm and 1050 nm, which provides insight into the spectral dispersion of the nanodiamond formulations. The two measured wavelengths provide a limited indication of dispersion; refractive index was not measured at the 1300-nm wavelength used for three-photon imaging.

The RI differences observed across nanodiamond formulations identify a range of optical properties in the tested samples but do not establish precise control of RI by formulation. Although effects such as nanodiamond agglomeration^44^ likely contribute to the variability seen in the older sample set, these results point to yet unexplored headroom for optimisation. Future work could examine whether improved control over dispersion, aggregation and composition yields reproducibly targeted refractive-index ranges. Surface modification offers an additional route for tailoring nanodiamond interactions with the surrounding matrix^45^.

The nanodiamond phantoms provide an optically heterogeneous environment that differs from the relatively homogeneous agarose-bead reference. We therefore examined how measured PSF widths changed with imaging depth. PSFs were recorded in the same nano 10% phantom near its surface (0–50 µm) and approximately 700 µm deeper (Figure 2c), with identical excitation power and microscope settings; imaging depth was the only experimental variable. Gaussian fitting of lateral and axial profiles yielded the FWHM values shown in Figure 2d,e. The measured PSFs broadened at greater depth, consistent with depth-dependent optical effects within the phantom.

### Stability of measured PSF widths across excitation powers

Accurate PSF measurements require the apparent width of the fluorescent emitter to remain stable across the acquisition conditions. Photobleaching resistance of fluorescent nanodiamonds has been reported previously^23,25^. Here, we assessed the measured PSF widths of fluorescent beads and nanodiamonds across a range of two-photon excitation powers. These measurements characterise the stability of the fitted PSF widths; they do not independently distinguish fluorophore saturation from detector saturation or directly quantify photobleaching.

Figure 3a shows xz images of fluorescent beads, whereas Figure 3b shows the corresponding images of nanodiamonds (nano 8%) acquired under two-photon excitation at increasing laser powers. Both materials were imaged using identical PMT settings. At higher excitation powers, the bead images showed signal saturation accompanied by apparent PSF broadening, whereas nanodiamond fluorescence profiles remained better defined over the tested range. Gaussian fitting of axial intensity profiles quantified the power-dependent changes in z-FWHM shown in Figure 3c.

**Figure 3.**
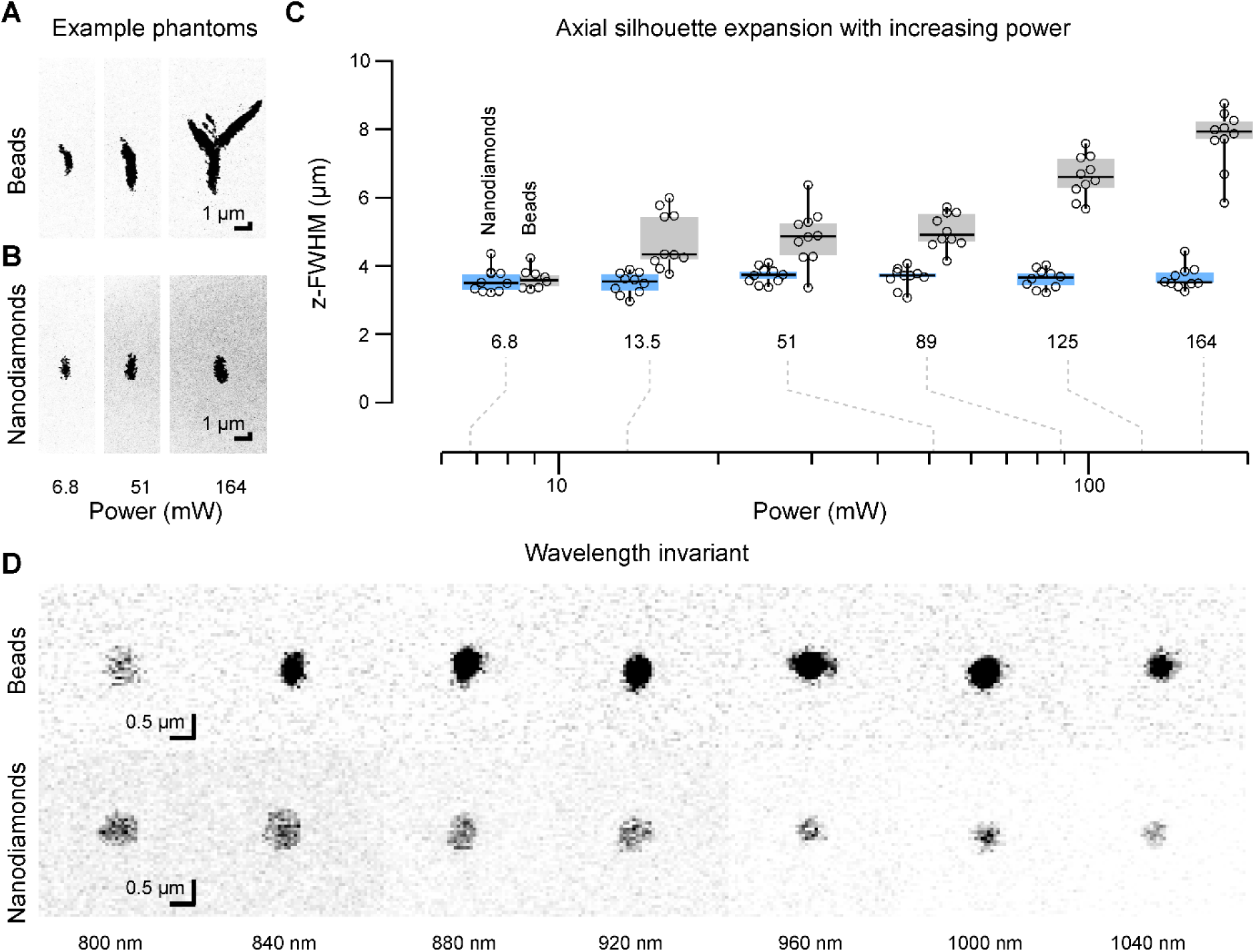
Power-dependent PSF widths and excitation-wavelength detectability under two-photon imaging. (a) Xz images of fluorescent beads at increasing two-photon excitation powers, acquired with the same PMT settings as for nanodiamonds. At higher powers, bead images show signal saturation and apparent PSF broadening; the origin of saturation was not independently established. (b) Corresponding xz images of nanodiamonds (nano 8% sample) showing well-preserved morphology across the same excitation power range, consistent with more stable measured PSF widths over the tested power range. (c) Axial z-FWHM values obtained from Gaussian fitting of the fluorescence profiles in (a,b). Nanodiamond-derived axial PSF widths remain comparatively stable across the tested power range, whereas bead-derived axial widths increase at higher power. This panel reports power-dependent axial PSF measurements, not bleaching-time measurements. Open circles indicate individual measurements; boxes show Q1–Q3 (Tukey quartiles), centre lines show the median, and whiskers span the mean ± 1 SD. (d) Fluorescence images of beads and nanodiamonds under two-photon excitation from 800 to 1040 nm at a constant average excitation power of 5 mW (Setup A), displayed without individual intensity normalisation. Chirp compensation and the optical configuration were not adjusted between wavelengths. Both materials remained detectable across the tested range; these data do not provide a quantitative two-photon excitation spectrum.

Across the tested excitation powers, nanodiamond-derived axial PSF widths remained comparatively stable, whereas bead-derived axial widths increased at higher powers (Figure 3c). The experiment quantified FWHM as a function of excitation power, not fluorescence intensity or bleaching over time. It does not independently establish whether the observed bead saturation arose in the fluorophore, detector or acquisition electronics.

### Nanodiamonds support robust imaging across a wide excitation spectrum

Figure 3d presents fluorescence images of beads and nanodiamonds acquired under two-photon excitation at wavelengths from 800 to 1040 nm and displayed without individual intensity normalisation. All measurements were acquired on Setup A using the same Coherent Vision S laser at a constant average excitation power of 5 mW, without changing the chirp compensation or microscope optical configuration between wavelengths. Both materials remained detectable throughout the tested range. Because pulse duration and dispersion were not characterised separately at each wavelength, these observations demonstrate practical detectability rather than a quantitative two-photon excitation spectrum.

### Nanodiamond PSF measurements are reproducible across microscope setups and over time

To test whether nanodiamond-based PSF benchmarking generalises beyond a single optical configuration, we compared PSF measurements from 8% dnp nanodiamonds and conventional fluorescent beads on three two-photon microscopes (Figure 4a,b). Within each setup, neither lateral nor axial PSF widths differed significantly between emitter types. However, the microscopes did not yield identical PSFs: Setup B showed broader axial profiles than Setup A with both reference materials. These findings support within-instrument benchmarking with nanodiamonds, although non-significant comparisons do not establish formal equivalence.

**Figure 4.**
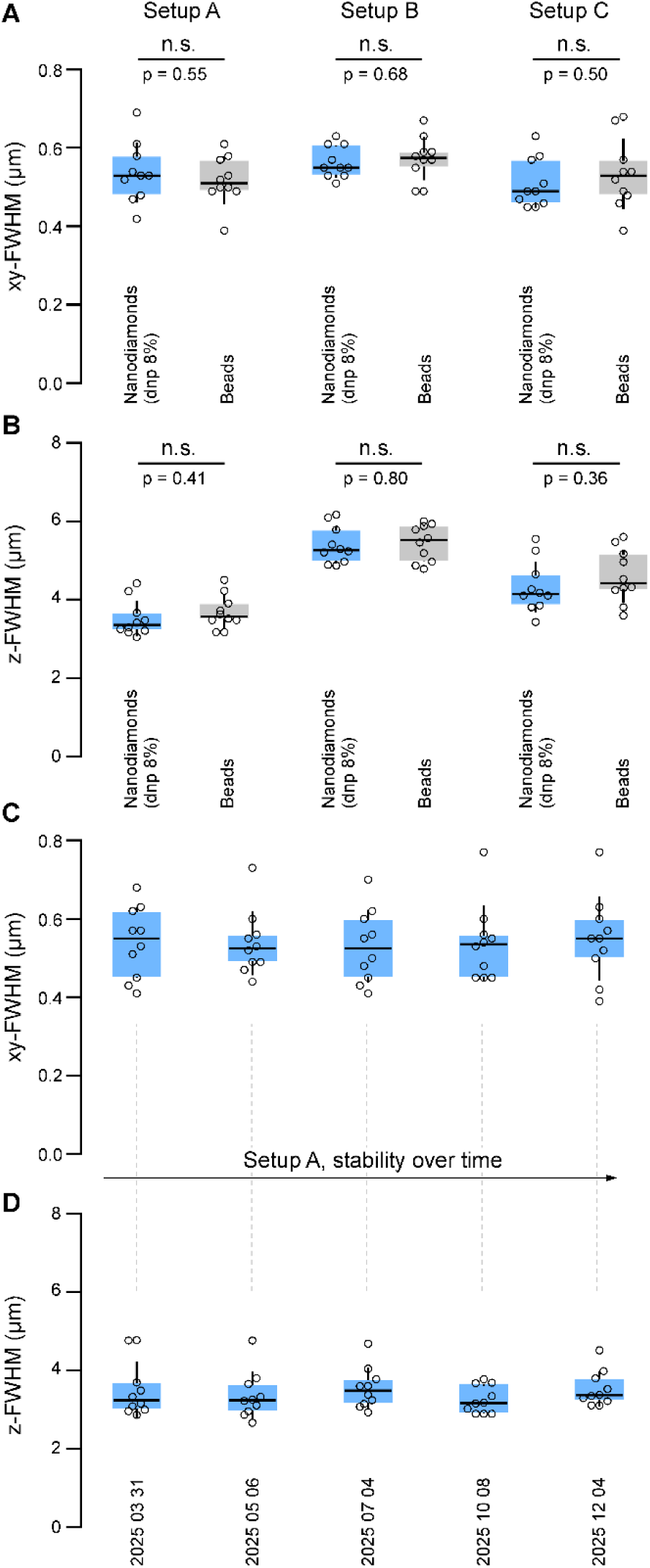
Cross-platform comparison and longitudinal stability of nanodiamond-based PSF measurements. (a) Lateral xy-FWHM values from dnp 8% nanodiamonds (blue) and fluorescent beads (grey) on three two-photon microscopes: A and B at the University of Sussex and C at the Francis Crick Institute. Mean ± SD (nanodiamonds versus beads; µm) was 0.54 ± 0.08 versus 0.52 ± 0.06 on A, 0.56 ± 0.04 versus 0.57 ± 0.06 on B, and 0.51 ± 0.06 versus 0.53 ± 0.09 on C. No within-setup difference was statistically significant (A: p = 0.55; B: p = 0.68; C: p = 0.50). (b) Corresponding axial z-FWHM measurements. Mean ± SD (nanodiamonds versus beads; µm) was 3.53 ± 0.46 versus 3.70 ± 0.43 on A, 5.40 ± 0.47 versus 5.45 ± 0.46 on B, and 4.33 ± 0.65 versus 4.61 ± 0.68 on C. No within-setup difference was statistically significant (A: p = 0.41; B: p = 0.80; C: p = 0.36). For (a,b), n = 10 isolated emitters per group; two-sided unpaired two-sample t-tests. Open circles indicate individual measurements; boxes extend from Q1 to Q3 (Tukey quartiles), centre lines indicate the median, and whiskers extend from the mean − 1 SD to the mean + 1 SD. (c,d) Repeated lateral and axial FWHM measurements of the dnp 8% nanodiamond phantom on Setup A across five sessions between March and December 2025. Open circles indicate individual emitters; the lower and upper edges of each box represent Q1 and Q3 (Tukey quartiles), its centre line indicates the median, and whiskers extend from the mean − 1 SD to the mean + 1 SD. The longitudinal measurements are presented descriptively.

We next assessed longitudinal stability using repeated measurements on Setup A (Figure 4c,d). Across five imaging sessions spanning March to December 2025, both lateral and axial nanodiamond-derived FWHM distributions remained tightly clustered without an obvious systematic drift. This stability supports the use of a long-lived nanodiamond phantom for longitudinal microscope quality control, where changes in measured PSF can be interpreted against a stable physical reference rather than a newly prepared sample.

### Nanodiamonds enable depth-aware PSF benchmarking under three-photon excitation

We next evaluated the nanodiamond phantoms under three-photon excitation at 1300 nm (Figure 5). At approximately 220 µm depth, nanodiamonds and beads yielded similar lateral and axial PSF widths at identical excitation power (4.85 mW); neither comparison reached statistical significance. These tests do not establish formal equivalence between the two emitter types. At approximately 720 µm depth and the same excitation power, nanodiamond-derived PSFs were broader than bead-derived PSFs in both dimensions (Figure 5b,c). Most notably, axial FWHM in the nanodiamond phantom increased from 1.50 ± 0.17 µm at approximately 220 µm to 2.33 ± 0.39 µm at approximately 720 µm, whereas the corresponding bead-reference measurements remained close to 1.6 µm. These differences are consistent with depth-dependent optical effects within the heterogeneous phantom, although differences in the population of detectable aggregates at depth cannot be excluded. Nanodiamond PSFs remained measurable at approximately 720 µm at the additional excitation power of 14.5 mW; this condition was not included in equal-power comparisons with beads.

**Figure 5.**
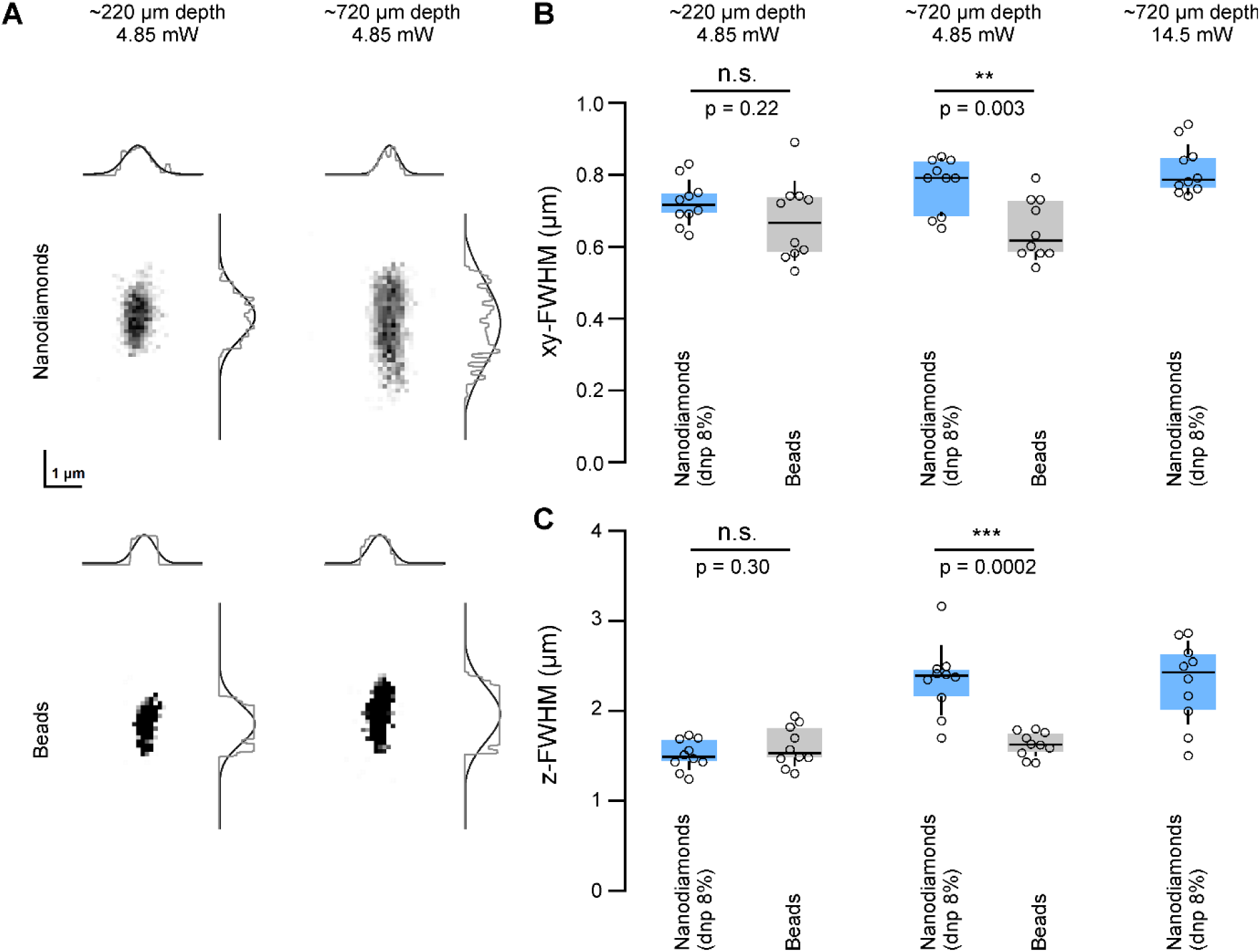
Three-photon PSF benchmarking with nanodiamond phantoms. (a) Representative three-photon PSFs and corresponding lateral/axial intensity profiles for 8% dnp nanodiamonds and fluorescent beads at approximately 220 µm and 720 µm imaging depth, acquired at 1300 nm and 4.85 mW. Scale bar, 1 µm. (b) Lateral xy-FWHM at approximately 220 and 720 µm imaging depth, at 4.85 mW. Mean ± SD (nanodiamonds versus beads; µm) was 0.72 ± 0.06 versus 0.67 ± 0.11 at approximately 220 µm (p = 0.22) and 0.77 ± 0.08 versus 0.65 ± 0.08 at approximately 720 µm (p = 0.003). The additional nanodiamond condition at approximately 720 µm and 14.5 mW yielded 0.81 ± 0.07 µm and was not included in the equal-power bead comparisons. (c) Axial z-FWHM at approximately 220 and 720 µm imaging depth, at 4.85 mW. Mean ± SD (nanodiamonds versus beads; µm) was 1.50 ± 0.17 versus 1.59 ± 0.22 at approximately 220 µm (p = 0.30) and 2.33 ± 0.39 versus 1.62 ± 0.14 at approximately 720 µm (p = 0.0002). At approximately 720 µm and 14.5 mW, nanodiamond z-FWHM was 2.31 ± 0.47 µm. For (b,c), n = 10 isolated emitters per condition; pairwise nanodiamond–bead comparisons used two-sided Welch’s t-tests. Open circles indicate individual measurements, and boxes summarise the distributions.

## Discussion

Reliable measurement of spatial resolution is important for both two- and three-photon microscopy. Here, we evaluated two related uses of fluorescent nanodiamond phantoms: reusable fluorescent emitters for repeated PSF measurement and an optically heterogeneous environment for assessing changes in the effective PSF with imaging depth^22^. In all three tested two-photon microscopes, no statistically significant nanodiamond–bead difference in lateral or axial FWHM was detected within each setup. The same phantom also supported repeated measurements across five sessions spanning March to December 2025.

The second application concerns depth-dependent imaging performance. The measured refractive indices of several nanodiamond formulations fall within ranges reported for some brain tissues, while the phantom exhibits greater visible heterogeneity than the agarose-bead reference. Under three-photon excitation at 4.85 mW, mean axial FWHM in the nanodiamond phantom increased from 1.50 to 2.33 µm (approximately 55%) between imaging depths of approximately 220 and 720 µm, whereas the bead reference yielded 1.59 and 1.62 µm at the respective depths (Figure 5). At approximately 720 µm, nanodiamond-derived axial PSFs were broader than bead-derived PSFs measured at the same excitation power. This observation is consistent with depth-dependent changes in focusing within the phantom, but the contributions of scattering, refractive-index mismatch and possible emitter-selection effects cannot be separated from these data alone. PSF characterisation in turbid phantoms provides an established framework for assessing how the surrounding medium affects two-photon imaging performance^46^.

The measured differences in axial PSF width also illustrate why calibration conditions matter for quantitative image interpretation. Under a Gaussian convolution model, using a narrower calibration PSF yields a larger inferred object width for the same measured image profile. However, the present experiments do not establish which phantom more accurately reproduces the PSF within biological tissue; this would require validation against an independent reference under matched imaging conditions.

For instrument-specific benchmarking, agreement between different fluorescent reference materials must be distinguished from agreement between microscope configurations. Setup B exhibited broader axial PSFs than Setup A for both beads and nanodiamonds (Figure 4b). A reusable phantom can therefore complement conventional bead-based references when following the same microscope over time, provided that imaging depth, acquisition settings and analysis procedures are reported consistently^22,47^.

The samples were originally fabricated in 2018, kept in small Petri dishes at room temperature, and used again in 2025 without visible signs of degradation during imaging. This demonstrates their continued practical usability after extended storage; direct baseline fluorescence measurements from 2018 were not available to quantify any changes over that period. Repeated PSF measurements of a stored phantom across five sessions in 2025 showed no obvious systematic drift, illustrating its potential for longitudinal instrument checks.

Several limitations define the scope of these conclusions. Non-significant bead–nanodiamond comparisons do not establish formal metrological equivalence, and the full size distribution of the fluorescent aggregates selected for PSF analysis was not independently determined. Scattering coefficients were not measured, and no matched two-photon depth series was acquired in the agarose-bead reference. Consequently, the observed broadening cannot be attributed uniquely to scattering or interpreted as a direct comparison of depth-dependent performance between the two materials. The refractive indices of the phantom were not assessed at the 1300-nm three-photon excitation wavelength, and possible depth-related differences in the population of detectable emitters cannot be excluded. The available fabrication history does not establish batch-to-batch reproducibility; the present longitudinal measurements concern the same stored phantom. Absolute lateral FWHM estimates are limited by spatial sampling, particularly on Setups A and C; identical within-setup acquisition settings for beads and nanodiamonds do not remove this limitation. The punctate confocal fluorescence was not verified using a matrix-only or emission-spectral control, and the power-series experiment does not provide a direct bleaching assay. Blinking has been reported for individual nitrogen-vacancy centres in discrete nanodiamonds^48^. The present measurements did not characterise emitter-level fluorescence intermittency.

An additional comparison of beads embedded in a matrix with matched optical properties would help distinguish the effects of emitter material from those of the surrounding medium. Such a control would refine the physical interpretation of depth-dependent differences, while the present data establish the feasibility of recording repeated nanodiamond-derived PSF measurements across microscope configurations and imaging depths.

Together, the results support fluorescent nanodiamond phantoms as durable, reusable materials for PSF benchmarking under the tested two- and three-photon imaging conditions. Their combination of repeated reference measurements and sensitivity to depth-dependent changes in the effective PSF offers a practical complement to conventional fluorescent-bead samples. Further control of aggregate properties and phantom composition could support more standardised reference materials for deep multiphoton microscopy.

## Methods

### RI measurements

The refractive index at 589 nm and 1050 nm was measured using an Atago Multi-Wavelength Abbe Refractometer (model DR-M2). To maintain uniformity and reduce thermal variability in the RI values, all measurements were carried out at a regulated temperature of 22 °C.

### Mouse brain photo

The photograph shows the exposed surface of an adult mouse brain (C57BL/6J genetic background) taken through a cranial window at the University of Sussex. The photos of the agarose sample and phantoms were taken by the same camera in the same illumination conditions. A circular craniotomy (3 mm diameter; coordinates from bregma: 1mm caudal, 3mm to the right) was performed to allow for accessing the barrel cortex (whisker-related area of the primary somatosensory cortex (wS1) in mice). The animal procedure associated with the mouse brain photograph in Figure 2a was conducted under UK Home Office Project Licence 70/8400, following review by the University of Sussex Animal Welfare and Ethical Review Body and the Home Office, in compliance with the UK Animals (Scientific Procedures) Act 1986 and European Directive 2010/63/EU.

### Scanning Electron Microscopy images

SEM imaging was carried out using an SM-7800F Schottky Field Emission Scanning Electron Microscope (JEOL Ltd., Japan). Multiple magnifications were applied to capture both micro- and nanoscale features of the samples. A low-voltage electron source (1.00 kV) was used to prevent surface melting of the agar–carrageenan matrix observed at higher beam energies. Acquired images were post-processed using ImageJ software (version 1.54g) in combination with the DeconvolutionLab2 plugin (version 2.1.2). The Richardson–Lucy deconvolution algorithm was applied with 10 iterations to enhance image clarity and contrast.

### Two-photon microscope setups

The two- and three-photon imaging systems used in this study are summarised in Supplementary Table S1.

Setup A (University of Sussex). The excitation beam was generated using a tuneable femtosecond Ti:Sapphire laser (Coherent Vision-S, 75 fs, 80 MHz, >2.5 W). The beam passed through an achromatic half-wave plate (AHWP05M-980, Thorlabs) and was split equally into two independent two-photon (2P) setups using a beam-splitter (10RQ00UB.4, Newport). It then passed through a Pockels cell (350-80, Conoptics), a telescope (AC254-075-B and AC254-150-B, Thorlabs), and was reflected by three silver mirrors (PF10-03-P01) into the head of a Sutter MOM stage. The beam was directed using galvanometric mirrors (6215H, Cambridge Technology). In the diffraction-limited configuration, it was focused into a 50 mm focal length scan lens (VISIR 1534SPR136, Leica) and further collimated by a 200 mm focal length tube lens (MXA22018, Nikon). In the Non-telecentric configuration (nTC1), the scan lens was modified to shift its focal length to 190 mm, and an additional plano-convex lens (175 mm) was added. The beam was reflected by two silver parabolic mirrors and passed through a dichroic mirror (T470/640rpc, Chroma) before slightly overfilling the back aperture of a Zeiss Objective W “Plan-Apochromat” ×20/1.0 to create a diffraction-limited excitation spot at a working distance of 1.8 mm. Fluorescence light was collected exclusively through the objective. A dichroic mirror (T470/640rpc, Chroma) reflected the fluorescence into a 140 mm focal length collecting lens, followed by a 580-nm dichroic mirror (H 568 LPXR, superflat). The signal was split into green and red channels using bandpass filters (ET525/50 and ET605/50, Chroma) and focused onto PMT detectors (H10770PA-40, Hamamatsu) using an aspheric condenser lens (G317703000, Linos). Images were acquired using custom-written ScanM software running in IGOR Pro 6.3 (WaveMetrics). Laser blanking occurred during the turnarounds using the Pockels cell.

Setup B (University of Sussex). Measurements were performed on a custom-built two-photon microscope based on a Sutter BOB platform (Sutter Instrument) equipped with a resonant–galvo–galvo (RGG) scanning system. Excitation was provided by a Coherent Axon 920 femtosecond laser operating at 920 nm. Imaging was performed using the same Zeiss W Plan-Apochromat 20×/1.0 water-immersion objective as in Setup A. Fluorescence was detected using gated H11526-20-NF photomultiplier tubes (Hamamatsu). Microscope control and image acquisition were performed using ScanImage.

Setup C (The Francis Crick Institute). Measurements were performed on a custom-built INSS two-photon microscope equipped with a resonant–galvo–galvo (RGG) scanning system. Excitation was provided by a Coherent Vision-S femtosecond Ti:Sapphire laser operating at 920 nm. Imaging was performed using the same Zeiss W Plan-Apochromat 20×/1.0 water-immersion objective as in Setup A. Fluorescence was collected using the INSS collection optics and detected using H11706P-40 photomultiplier tubes (Hamamatsu). The emission filters were ET525/50 and ET605/50 (Chroma), matching those used in Setup A. Microscope control and image acquisition were performed using ScanImage.

For the cross-platform comparisons in Figure 4, nanodiamonds and fluorescent beads were imaged using identical acquisition settings within each microscope. On Setup A, images were acquired at 256 × 256 pixels over a 150 × 150 µm field of view (lateral pixel size, 0.586 × 0.586 µm), with a 0.2 µm axial step and 3 mW excitation power at the sample. On Setup B, images were acquired at 512 × 512 pixels over a 196 × 196 µm field of view (lateral pixel size, 0.383 × 0.383 µm), with a 0.2 µm axial step and 7.8 mW at the sample. On Setup C, images were acquired at 512 × 512 pixels over a 312 × 312 µm field of view (lateral pixel size, 0.609 × 0.609 µm), with a 0.2 µm axial step and 3 mW at the sample.

### Three-photon imaging

Three-photon PSF measurements were acquired from 8% dnp nanodiamond phantoms and conventional fluorescent-bead samples at imaging depths of approximately 220 µm and 720 µm. Pairwise nanodiamond-bead measurements shown in Figure 5 used 4.85 mW excitation power, and an additional nanodiamond condition at approximately 720 µm was acquired at 14.5 mW. Three-dimensional image stacks of isolated fluorescent emitters were used to quantify lateral (xy) and axial (z) PSF widths.

Three-photon imaging was performed using a custom-built microscope based on a Thorlabs Bergamo multiphoton platform equipped with a wavelength-tunable excitation source (Spirit-NOPA, Spectra-Physics; 1200–2500 nm) pumped by a Spirit-1030-100 femtosecond laser (Spectra-Physics). For the experiments reported here, excitation was performed at 1300 nm with a repetition rate of 2 MHz and a pulse duration of approximately 50 fs. Dispersion compensation was controlled using a prism-based FemtoControl system (APE). Imaging was performed with a 25× water-immersion objective (XLPLN25XWMP2, Olympus; NA 1.05). Fluorescence was collected in non-descanned mode using two GaAsP photomultiplier tubes (PMT2100, Thorlabs) mounted in a 14° detector module (BDM3214-3P, Thorlabs). Nanodiamond fluorescence was separated using a 562 nm dichroic mirror (FF562-Di03) and detected through a 525/50 nm bandpass filter (FF03-525/50, Semrock). Image acquisition was controlled using ThorImage 4.4, with xyz positioning provided by an MCM3000 motorized stage (Thorlabs). The microscope setup has been described previously^49^. All three-photon image stacks were acquired using the same acquisition settings: 512 × 512 pixels, lateral pixel sampling of 0.406 × 0.406 µm, an axial step size of 0.2 µm (201 optical sections per stack), a pixel dwell time of 1.98 µs and a frame rate of approximately 1.412 Hz. The excitation powers of 4.85 and 14.5 mW were measured below the objective.

### PSF fitting and statistical analysis

For each experimental condition, three-dimensional image stacks were visually inspected in a 3D preview. Ten fluorescent emitters were chosen by visual inspection from different optical sections, without a predefined spatial location. Emitters were considered suitable when sufficiently isolated, with a well-defined fluorescence profile and adequate signal for fitting; overlapping, markedly irregular or very dim structures were excluded. Lateral (xy) and axial (z) intensity profiles were fitted with Gaussian functions incorporating a constant background offset. FWHM was calculated from the fitted Gaussian standard deviation (σ) as 2√(2ln2)σ. An emitter was excluded after selection only if a meaningful Gaussian fit could not be obtained; minor deviations from a Gaussian profile were not used as an exclusion criterion. The same visual selection, fitting and exclusion procedure was applied to nanodiamonds and fluorescent beads on all microscopes. Selection was visually guided, not formal random sampling.

For the nanodiamond-versus-bead comparisons on the three two-photon microscopes shown in Figure 4a,b, n = 10 isolated emitters were analysed per group using two-sided unpaired two-sample t-tests (ttest2, MATLAB). Repeated nanodiamond measurements in Figure 4c,d are presented descriptively to assess stability of the measured PSF distributions over time; no inferential longitudinal test was performed.

For the pairwise nanodiamond-versus-bead comparisons in Figure 5, n = 10 isolated emitters were analysed per group and two-sided Welch’s t-tests were used. Exact p values are reported in the figure and legend. Statistical significance was assessed at p < 0.05. Non-significant differences were not interpreted as evidence of formal equivalence.

For the comparisons of nanodiamonds and fluorescent beads in the diffraction-limited and non-telecentric configurations (Figure 1d), n = 10 isolated emitters per group were analysed using two-sided unpaired two-sample t-tests (MATLAB ttest2). The six nanodiamond formulations in Figure 1e were compared by one-way ANOVA, separately for lateral and axial FWHM (n = 10 emitters per formulation). Depth-dependent lateral and axial FWHM measurements in Figure 2d,e were assessed separately using one-way ANOVA (n = 10 emitters per depth); the respective p values shown in the figure are 0.014 and 0.009. Measurements at the six excitation powers in Figure 3c (n = 10 emitters per material and power) are presented descriptively. No post-hoc pairwise comparisons are reported for the omnibus ANOVAs in Figure 1e, and no multiplicity correction was applied to the reported pairwise t-tests.

### Fabrication of agarose-based sample with beads

We utilized 0.175 ± 0.005 µm yellow-green (505/515) fluorescent beads (P7220, Invitrogen) for the calibration and imaging experiments. These beads were specifically chosen for their consistent size and well-characterized fluorescent properties, which are ideal for evaluating the resolution and point spread function (PSF) of the optical system. To prepare the calibration samples, the beads were carefully embedded within a block of low melting point agarose (Fisher Scientific, BP1360-100) at a depth of 1 mm. The agarose concentration in the bead-containing PSF reference was 0.75%, ensuring an optimal balance between sufficient rigidity for immobilizing the beads and minimal optical scattering to maintain clarity during imaging. This arrangement provided a stable and reproducible medium that allowed for accurate imaging and quantification of the optical properties, making it a practical reference for PSF measurements. The separately prepared 2% agarose sample plotted in Figure 2b was used for refractive-index comparison, not as the bead-containing PSF reference.

### Fabrication of nanodiamond phantoms

Nanodiamond-based phantoms were fabricated according to the previously published agar–carrageenan protocol (ref. 28). A gel base containing 1.413 g agar and 2.0 g carrageenan per 100 mL distilled water was heated to boiling, mixed and autoclaved for approximately 45 min at 126 °C. The hot gel base was then added to the prepared nanodiamond suspensions, mixed thoroughly and subjected to a second autoclaving step before the mixtures were poured into moulds, covered and allowed to solidify. The nano series contained 8%, 10% and 12% (v/v) of an ND-standard suspension in DMSO (Adámas Nanotechnologies; nominal particle size, 5 nm). The dnp series contained the corresponding proportions of a hydrophilic RT-DND-L nanodiamond suspension (Ray Techniques Ltd.; nominal particle size, 4–5 nm), prepared as a 0.1% (w/w) dispersion in DMSO (ref. 28). The phantoms used in the present study were fabricated in Gdańsk in 2018 for tissue-mimicking material and MRI calibration studies. The exact preparation dates of individual formulations within this 2018 sample set were not recorded for the present optical study, and independent fabrication batches were not compared. The phantoms were stored at room temperature in small Petri dishes and reused for imaging in 2025. No visible drying, cracking or structural deterioration was observed during these experiments. Comparable fluorescence measurements from 2018 were not available to quantify changes in brightness or aggregate distribution during storage.

### Confocal images

Confocal images of samples were taken on a Leica TCS SP8 inverted microscope, using a HC PL APO 63x/1.20 water objective. Approximately 500 µm thick samples were coverslipped in aqua-poly/mount (Polysciences, 18606) and imaged with the 561 nm (69% laser power) laser line. Image stacks of 50×50×50 μm were acquired at a pixel size of 0.1 x 0.1 x 0.36 μm (x, y, z) and visualized as a 3D volume using napari. Fluorescence was collected using a broad detection window to maximize signal-to-noise in thick samples; no spectral unmixing or charge-state discrimination was performed.

## Data availability

The data supporting the findings of this study are available from the corresponding author upon reasonable request.

## Code availability

The MATLAB scripts used for PSF analysis and figure preparation are available from the corresponding author upon reasonable request.

## Acknowledgements

We thank Chrysovalantis Fekos and Miguel Maravall for the mouse brain photo used in Figure 2. Funding was provided by the Wellcome Trust (Investigator Award in Science 220277/Z20/Z), the European Research Council (ERC-StG “NeuroVisEco” 677687 and ERC AdG “Cones4Action” covered under the UK’s EPSRC guarantee scheme EP/Z533981/1), UKRI (BBSRC, BB/R014817/1 and BB/W013509/1), HFSP (RGP001/2025), the Leverhulme Trust (PLP-2017-005, RPG-2021-026 and RPG-2-23-042), the Lister Institute for Preventive Medicine and Gdańsk University of Technology by the DEC-4/1/2024/IDUB/I.1a/No grant under the NOBELIUM and 1/1/2025/IDUB/I.1B/Pt PLATINUM - ‘Excellence Initiative - Research University’ program. To promote Open Access, the authors have applied a CC BY public copyright licence to any Author Accepted Manuscript version arising from this submission. The authors declare no conflict of interest.

## Author contributions

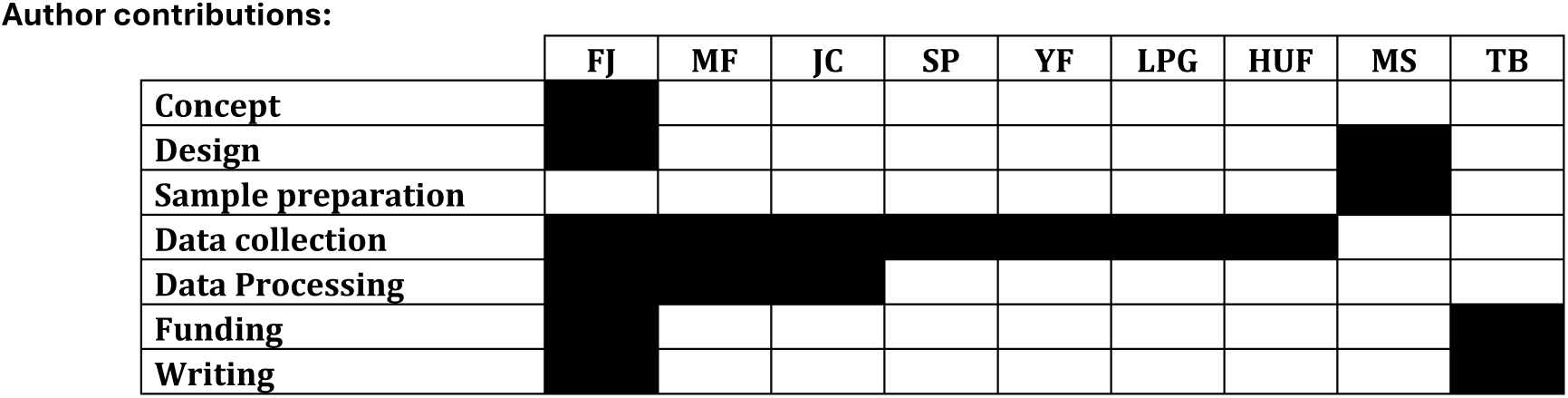

---

**Supplementary Table S1.**
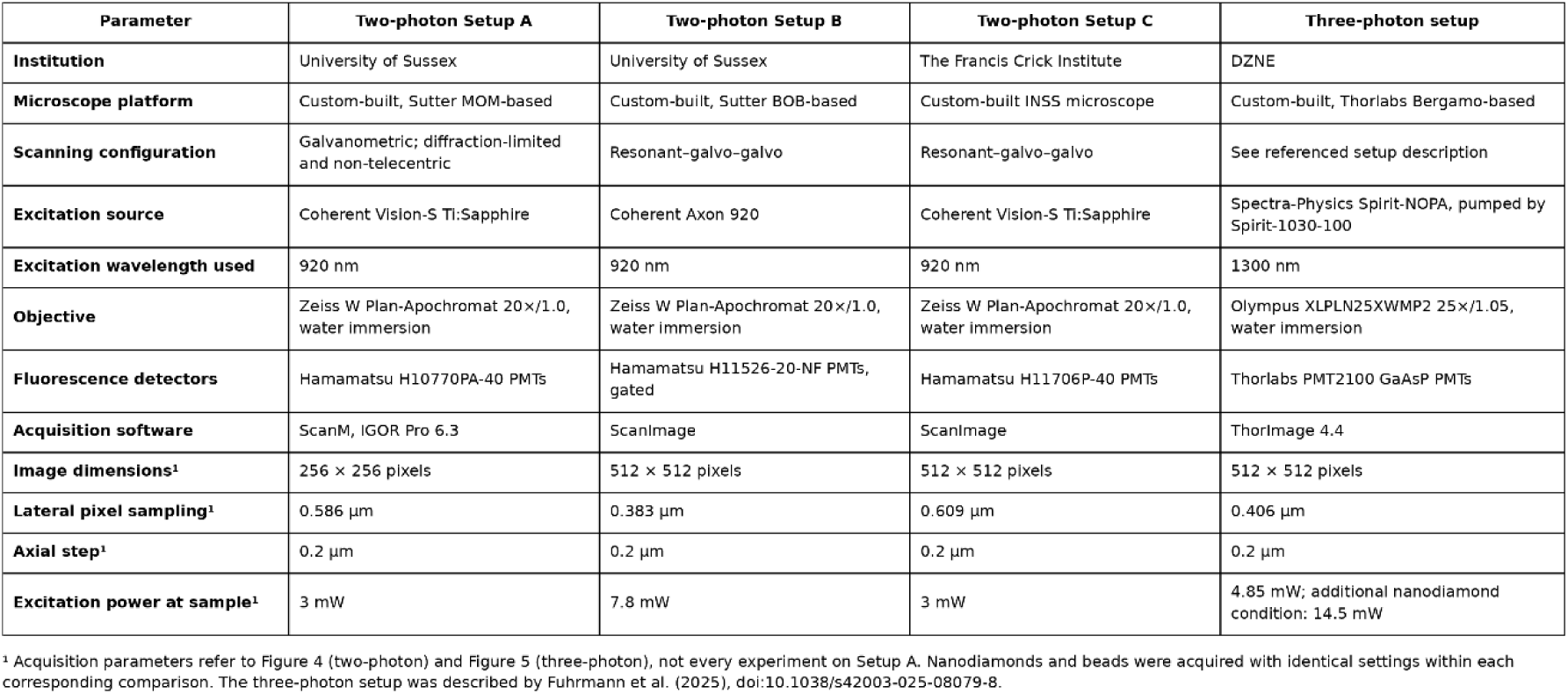
Microscope systems used for two- and three-photon PSF measurements. The table summarises acquisition parameters for Figures 4 and 5.

| Parameter | Two-photon Setup A | Two-photon Setup B | Two-photon Setup C | Three-photon setup |
| --- | --- | --- | --- | --- |
| Institution | University of Sussex | University of Sussex | The Francis Crick Institute | DZNE |
| Microscope platform | Custom-built, Sutter MOM-based | Custom-built, Sutter BOB-based | Custom-built INSS microscope | Custom-built, Thorlabs Bergamo-based |
| Scanning configuration | Galvanometric; diffraction-limited and non-telecentric | Resonant-galvo-galvo | Resonant-galvo-galvo | See referenced setup description |
| Excitation source | Coherent Vision-S Ti:Sapphire | Coherent Axon 920 | Coherent Vision-S Ti:Sapphire | Spectra-Physics Spirit-NOPA, pumped by Spirit-1030-100 |
| Excitation wavelength used | 920 nm | 920 nm | 920 nm | 1300 nm |
| Objective | Zeiss W Plan-Apochromat 20×/1.0, water immersion | Zeiss W Plan-Apochromat 20×/1.0, water immersion | Zeiss W Plan-Apochromat 20×/1.0, water immersion | Olympus XLPLN25XWMP2 25×/1.05, water immersion |
| Fluorescence detectors | Hamamatsu H10770PA-40 PMTs | Hamamatsu H11526-20-NF PMTs, gated | Hamamatsu H11706P-40 PMTs | Thorlabs PMT2100 GaAsP PMTs |
| Acquisition software | ScanM, IGOR Pro 6.3 | ScanImage | ScanImage | ThorImage 4.4 |
| Image dimensions<sup>1</sup> | 256 × 256 pixels | 512 × 512 pixels | 512 × 512 pixels | 512 × 512 pixels |
| Lateral pixel sampling<sup>1</sup> | 0.586 μm | 0.383 μm | 0.609 μm | 0.406 μm |
| Axial step<sup>1</sup> | 0.2 μm | 0.2 μm | 0.2 μm | 0.2 μm |
| Excitation power at sample<sup>1</sup> | 3 mW | 7.8 mW | 3 mW | 4.85 mW; additional nanodiamond condition: 14.5 mW |

<sup>1</sup> Acquisition parameters refer to Figure 4 (two-photon) and Figure 5 (three-photon), not every experiment on Setup A. Nanodiamonds and beads were acquired with identical settings within each corresponding comparison. The three-photon setup was described by Fuhrmann et al. (2025), doi:10.1038/s42003-025-08079-8.

## References

1. Denk, W., Strickler, J. H. & Webb, W. W. Two-Photon Laser Scanning Fluorescence Microscopy. Science 248, 73–76 (1990).

2. Zipfel, W. R., Williams, R. M. & Webb, W. W. Nonlinear magic: multiphoton microscopy in the biosciences. Nat. Biotechnol. 21, 1369–1377 (2003).

3. Svoboda, K. & Yasuda, R. Principles of Two-Photon Excitation Microscopy and Its Applications to Neuroscience. Neuron 50, 823–839 (2006).

4. Zhou, M. et al. Zebrafish Retinal Ganglion Cells Asymmetrically Encode Spectral and Temporal Information across Visual Space. Curr. Biol. 30, 2927–2942.e7 (2020).

5. Bando, Y., Wenzel, M. & Yuste, R. Simultaneous two-photon imaging of action potentials and subthreshold inputs in vivo. Nat. Commun. 12, 7229 (2021).

6. Dunn, T. W. et al. Neural Circuits Underlying Visually Evoked Escapes in Larval Zebrafish. Neuron 89, 613–628 (2016).

7. Ebina, T. et al. Two-photon imaging of neuronal activity in motor cortex of marmosets during upper-limb movement tasks. Nat. Commun. 9, 1879 (2018).

8. Bartel, P., Janiak, F. K., Osorio, D. & Baden, T. Colourfulness as a possible measure of object proximity in the larval zebrafish brain. Curr. Biol. 31, R235–R236 (2021).

9. Lees, R. M. et al. Standardized measurements for monitoring and comparing multiphoton microscope systems. Nat. Protoc. 1–38 (2025) doi:10.1038/s41596-024-01120-w.

10. Cole, R. W., Jinadasa, T. & Brown, C. M. Measuring and interpreting point spread functions to determine confocal microscope resolution and ensure quality control. Nat. Protoc. 6, 1929–1941 (2011).

11. Juškaitis, R. Measuring the Real Point Spread Function of High Numerical Aperture Microscope Objective Lenses. in Handbook Of Biological Confocal Microscopy (ed. Pawley, J. B.) 239–250 (Springer US, Boston, MA, 2006). doi:10.1007/978-0-387-45524-2_11.

12. Sofroniew, N. J., Flickinger, D., King, J. & Svoboda, K. A large field of view two-photon mesoscope with subcellular resolution for in vivo imaging. eLife 5, e14472 (2016).

13. North, A. J. Seeing is believing? A beginners’ guide to practical pitfalls in image acquisition. J. Cell Biol. 172, 9–18 (2006).

14. Egner, A. & Hell, S. W. Aberrations in Confocal and Multi-Photon Fluorescence Microscopy Induced by Refractive Index Mismatch. in Handbook Of Biological Confocal Microscopy (ed. Pawley, J. B.) 404–413 (Springer US, Boston, MA, 2006). doi:10.1007/978-0-387-45524-2_20.

15. Confocal and Two-Photon Microscopy: Foundations, Applications and Advances | Wiley. *Wiley.com* https://www.wiley.com/en-us/Confocal+and+Two-Photon+Microscopy%3A+Foundations%2C+Applications+and+Advances-p-9780471409205.

16. Horton, N. G. et al. In vivo three-photon microscopy of subcortical structures within an intact mouse brain. Nat. Photonics 7, 205–209 (2013).

17. Ouzounov, D. G. et al. In vivo three-photon imaging of activity of GCaMP6-labeled neurons deep in intact mouse brain. Nat. Methods 14, 388–390 (2017).

18. Prevedel, R. et al. Three-photon microscopy: an emerging technique for deep intravital brain imaging. Nat. Rev. Neurosci. 26, 521–537 (2025).

19. Yildirim, M., Sugihara, H., So, P. T. C. & Sur, M. Functional imaging of visual cortical layers and subplate in awake mice with optimized three-photon microscopy. Nat. Commun. 10, 177 (2019).

20. Streich, L. et al. High-resolution structural and functional deep brain imaging using adaptive optics three-photon microscopy. Nat. Methods 18, 1253–1258 (2021).

21. Faklaris, O. et al. Detection of Single Photoluminescent Diamond Nanoparticles in Cells and Study of the Internalization Pathway. Small 4, 2236–2239 (2008).

22. Žurauskas, M., Alex, A., Park, J., Hood, S. R. & Boppart, S. A. Fluorescent nanodiamonds for characterization of nonlinear microscopy systems. Photonics Res. 9, 2309–2318 (2021).

23. Tisler, J. et al. Fluorescence and Spin Properties of Defects in Single Digit Nanodiamonds. ACS Nano 3, 1959–1965 (2009).

24. Vaijayanthimala, V. et al. The long-term stability and biocompatibility of fluorescent nanodiamond as an *in vivo* contrast agent. Biomaterials 33, 7794–7802 (2012).

25. Alkahtani, M. H. et al. Fluorescent nanodiamonds: past, present, and future. Nanophotonics 7, 1423–1453 (2018).

26. Chauhan, S., Jain, N. & Nagaich, U. Nanodiamonds with powerful ability for drug delivery and biomedical applications: Recent updates on in vivo study and patents. J. Pharm. Anal. 10, 1–12 (2020).

27. Mochalin, V. N., Shenderova, O., Ho, D. & Gogotsi, Y. The properties and applications of nanodiamonds. Nat. Nanotechnol. 7, 11–23 (2012).

28. Sękowska, A. et al. Nanodiamond phantoms mimicking human liver: perspective to calibration of T1 relaxation time in magnetic resonance imaging. Sci. Rep. 10, 6446 (2020).

29. Shenderova, O., Hens, S. & McGuire, G. Seeding slurries based on detonation nanodiamond in DMSO. Diam. Relat. Mater. 19, 260–267 (2010).

30. Janiak, F. K. et al. Non-telecentric two-photon microscopy for 3D random access mesoscale imaging. Nat. Commun. 13, 544 (2022).

31. Abookasis, D. & Meitav, O. Assessing mouse brain tissue refractive index in the NIR spectral range utilizing spatial frequency domain imaging technique combined with processing algorithms. in vol. 10864 108640R (2019).

32. Lee, A. J. et al. Volumetric Refractive Index Measurement and Quantitative Density Analysis of Mouse Brain Tissue with Sub-Micrometer Spatial Resolution. Adv. Photonics Res. 4, 2300112 (2023).

33. Binding, J. et al. Brain refractive index measured in vivo with high-NA defocus-corrected full-field OCT and consequences for two-photon microscopy. Opt. Express 19, 4833–4847 (2011).

34. Ryu, Y., et al. Single-Step Fast Tissue Clearing of Thick Mouse Brain Tissue for Multi-Dimensional High-Resolution Imaging. Int. J. Mol. Sci. 23, 6826 (2022).

35. Sun, J., Lee, S. J., Wu, L., Sarntinoranont, M. & Xie, H. Refractive index measurement of acute rat brain tissue slices using optical coherence tomography. Opt. Express 20, 1084–1095 (2012).

36. Schlüßler, R. et al. Mechanical Mapping of Spinal Cord Growth and Repair in Living Zebrafish Larvae by Brillouin Imaging. Biophys. J. 115, 911–923 (2018).

37. Boothe, T. et al. A tunable refractive index matching medium for live imaging cells, tissues and model organisms. eLife https://elifesciences.org/articles/27240 (2017) doi:10.7554/eLife.27240.

38. Philipp, K. et al. Diffraction-limited axial scanning in thick biological tissue with an aberration-correcting adaptive lens. Sci. Rep. 9, 9532 (2019).

39. Horst, J. van der, Trull, A. K. & Kalkman, J. Deep-tissue label-free quantitative optical tomography. Optica 7, 1682–1689 (2020).

40. Favre-Bulle, I. A. et al. Scattering of Sculpted Light in Intact Brain Tissue, with implications for Optogenetics. Sci. Rep. 5, 11501 (2015).

41. Pende, M. et al. High-resolution ultramicroscopy of the developing and adult nervous system in optically cleared Drosophila melanogaster. Nat. Commun. 9, 4731 (2018).

42. Bekkouche, B. M. B., Fritz, H. K. M., Rigosi, E. & O’Carroll, D. C. Comparison of Transparency and Shrinkage During Clearing of Insect Brains Using Media With Tunable Refractive Index. Front. Neuroanat. 14, (2020).

43. Ke, M.-T. et al. Super-Resolution Mapping of Neuronal Circuitry With an Index-Optimized Clearing Agent. Cell Rep. 14, 2718–2732 (2016).

44. Voitylov, A. V. et al. Light refraction in aqueous suspensions of diamond particles. Colloids Surf. Physicochem. Eng. Asp. 538, 417–422 (2018).

45. Krueger, A. & Lang, D. Functionality is Key: Recent Progress in the Surface Modification of Nanodiamond. Adv. Funct. Mater. 22, 890–906 (2012).

46. Dong, C.-Y., Koenig, K. & So, P. Characterizing point spread functions of two-photon fluorescence microscopy in turbid medium. J. Biomed. Opt. 8, 450 (2003).

47. Nelson, G. et al. QUAREP-LiMi: A community-driven initiative to establish guidelines for quality assessment and reproducibility for instruments and images in light microscopy. J. Microsc. 284, 56–73 (2021).

48. Bradac, C. et al. Observation and control of blinking nitrogen-vacancy centres in discrete nanodiamonds. Nat. Nanotechnol. 5, 345–349 (2010).

49. Fuhrmann, F. et al. Three-photon in vivo imaging of neurons and glia in the medial prefrontal cortex with sub-cellular resolution. *Commun*. Biol. 8, 795 (2025).

